# Behaviour of two species of psittacine birds at wild bird feeding sites in Australia

**DOI:** 10.1101/2020.10.27.356865

**Authors:** Michelle R. Plant, Dianne Vankan, Gregory Baxter, Evelyn Hall, David Phalen

## Abstract

This study investigated how Crimson Rosellas *Platycercus elegans* (CR) and Australian King-Parrots *Alisterus scapularis* (AKP) used provisioned seed at two public bird feeding sites in Australia. A total of 197 CR and 72 AKP were trapped and colour-banded. Observational data was collected every 10mins between 08:00-16:00 for three consecutive days during autumn and spring. Foraging effort was described using five metrics that quantified the birds’ visiting frequency and foraging duration over each day and observation period. Seed selection (over 5mins) and intake (over 10mins) were determined, and the energy intake was calculated. Total counts and population estimates were calculated for each species. Individual, species, seasonal and geographic variation in the use of provisioned seed was demonstrated by the metric summaries and Restricted Maximum Likelihood Modelling. Both species fed as part of large mixed species flocks that would not naturally congregate together to forage. Overall, CR were found to have higher foraging effort and feed in greater numbers than AKP, but a spectrum of use was observed for both species. Individuals were observed using the provisioned seed between 0-3 days/observation period. When birds used the provisioned seed, they were found to make between 1-8 visits/day, with most lasting 10-30mins. Few daily durations lasted longer than 50mins. Within a 10-minute interval, it was possible for a CR and AKP to obtain between 1.73-62.91% and 6.84-88.54% of their daily energy requirements, respectively. In a visit, either species could fill their crop and meet most, if not all, of their daily energy requirements. A small percentage of birds (6.5%) were found to use the feeding sites daily and for long durations (up to 160mins). It is likely that a proportion of the birds using the provisioned seed at both sites were dependent on the food source and would be at risk if the seed supply were suddenly reduced. The study also provided evidence that wild bird feeding provided an advantage to one or more species, as well as evidence that the food source did not affect the study species’ seasonal dispersal patterns or juveniles’ ability to forage on natural food sources.

## 1. Introduction

Wild bird feeding is one of the most common forms of wildlife interaction in the world (1-4). It is particularly popular in developed countries with birds being fed at up to 75% of people’s homes (2, 5-12). The primary reasons people engage in wild bird feeding are to experience the pleasure the activity provides and to compensate for habitat modifcation or harsh climatic conditions (6, 9, 13-15). Despite its popularity, wild bird feeding has elicited extensive criticism (16-19). Authors argue that wild bird feeding activities threaten to alter natural foraging behaviour, resulting in habitutation (17), reduced vigilance (16), impaired foraging skills (16, 20, 21), dependency on the suppled food (4, 16, 17, 19, 22), distruption of daily activity patterns (16, 17) and changes in species movement patterns (16, 20). Wild bird feeding may advantage dominant individuals or species (5, 9, 16, 23) and provide support to invasive species (24-27). There are also health concerns that include, but are not limited to, increased disease transmission between birds congregating at fixed point feeding sites and the spread of zoonotic diseases to humans (27, 28). Individually, or as a result of their interactions, these impacts could then contribute to carry-over effects on recruitment, reproduction and survival (29, 30), ultimately disrupting natural selection processes (22), particularly if the foraging resources provide support for less competitive individuals (30). Despite these often-raised concerns, empircal in situ data to validate them has been limited (18).

Authors have recommended that wild bird feeding studies investigating these concerns should begin by assessing the birds’ use of the provisioned foraging resources (31). To date, research on foraging effort at wild bird feeding sites has been dominated by studies of passerines in the United Kingdom and the United States of America (1, 30, 32-37), and a small number of studies have quantified the foraging effort of carnivorous species (38) and waterbirds (39). Foraging effort has been found to be influneced by the species studied (8, 23, 36), time of day (1, 33, 34, 38), location and the number of feeding opportunities within a bird’s home range (1, 34, 36). Differences in foraging effort have also been found to be impacted by the effects of temperature (1, 33, 34), season (8, 23, 34), individual bird personality (36, 37) and demographics—such as age and gender (8, 30, 34-36). Added complexities to assessing the factors impacting foraging effort include dominance hierarchies, geographical location (1, 31), and the feed type and it’s presentation (32, 40). Therefore, new studies should be conducted over multiple seasons (20) and locations (22), extend to a broader range of species being fed, and where possible, be conducted under standard feeding practices in the settings where the activity occurs (20).

In Australia, wild bird feeding takes place in a variety of settings including backyards (6, 41), public recreation areas (19, 39) and as part of wildlife tourism activities, which are often co-located with protected areas (42, 43). Australians enjoy feeding a wide variety of avifauna (19, 39), including a range of endemic parrots and cockatoos (Psittaciformes) (41, 44, 45). These birds are common and abundant visitors to backyard feeding sites (46-48) and the primary species targeted by wildlife tourism activities (42, 49, 50). Due to their numbers, Psittaciformes are likely the most commonly fed birds in Australia. However, to date, the only research on how native free-ranging Psittaciformes use human provisioned supplemental feed and the impacts that feeding may have on them has pertained to the Rainbow Lorikeet (*Trichoglossus moluccanus*) and the Scaly-breasted Lorikeet (*Trichoglossus chlorolepidotus*) (49).

This study aimed to investigate the validity of published concerns about the behavioural impacts of wild bird feeding for two psittacine species—Crimson Rosella (*Platycercus elegans*) and Australian King-Parrot (*Alisterus scapularis)*—by quantifying birds’ foraging efforts during autumn and spring at two geographically distinct public feeding sites in eastern Australia.

## 2. Methods

### 2.1 Study sites

The study was conducted at two sites, located approximately 1,300km apart, on the east coast of Australia. The first (site 1), was in Queensland (S 28°13’50.10”, E 153°08’07.54”, 915m) at O’Reilly’s Rainforest Retreat immediately adjacent to Lamington National Park, which is part of Gondwana Rainforests, a 370,000-hectare subtropical World Heritage listed protected area. Birds have been fed here since the 1930s (pers. comm. S. O’Reilly 09 Aug, 19). The second location was in Victoria where the primary site (site 2a) was at Grants Picnic Ground (S 37°53’13”, E 145°22’13”, 339m). Additional birds were captured and monitored at Sherbrooke Picnic Ground (site 2b; S 37°52’52”, E 145°21’34”, 487m) located approximately 1.2km from 2a. Both 2a and 2b were on public land associated with the Dandenong Ranges National Park, a 3,540-hectare protected area dominated by temperate forest. Compared to the area surrounding site 1, the Dandenong Ranges National Park had a higher level of fragmentation by built infrastructure—with dwellings and other businesses offering alternate bird feeding opportunities. While a semi-organised wild bird feeding activity had only been in operation at site 2a since 1999, birds were previously fed in an ad hoc way for an additional 40 years prior to this (pers. comm. M. Hoogland 09 Aug, 19). Regular opportunistic wild bird feeding also took place at site 2b, but the timeframe was not known.

### 2.2 Foraging resource

Operators at both feeding sites sold small packets of seed (primarily millet) to visitors who were able to hand feed birds within designated feeding areas. The site operators did not, themselves, provide feed to the birds. Regularly, some visitors brought their own feed including wild bird mix, black and grey-striped sunflower seeds and sometimes human food products.

### 2.3 Study population

The species studied at both sites were Crimson Rosellas (length 35-38cm, weight 120-150g) and Australian King-Parrots (length 40-45cm, weight 210-275g) (distribution see (51); description see (52, 53)). Of the species observed foraging at the study sites, these two were selected as they were common to both locations. Both species make use of similar forest and woodland habitats throughout their ranges in eastern and southeastern Australia (52). Resident birds remain within their territories (often at higher altitudes) year round, and non-breeding adults and sub-adults are known to dispese to lower lying areas during autumn and winter (44, 52). Both species mostly forage as pairs or small groups in the canopy, shrubs or at the ground, consuming plant material from native and introduced species, as well as insects (52, 54).

Birds were captured according to a randomised, stratified sampling methodology. Each feeding site was stratified into sections (Qld: north, south, east, west; Vic: 2a-1, 2a-2, 2a-3, 2b) and points where birds accessed the seed (hand/ground). A section and a feeding point were randomly drawn prior to each catching effort. If a bird was not available at the pre-selected point, then the closest bird was caught. Birds were caught using hand nets or drop cages (Professional Trapping Supplies, Molendinar, Qld, Aust.), and then transferred to a calico bag until sampled and colour-banded.

In Queensland, bird capture and banding was conducted over 19 days between spring 2007 and winter 2008. Birds were colour-banded using the Australian Bird and Bat Banding Scheme’s schema 02 (55). Coloured bands were obtained from A.C. Hughes, United Kingdom and Lentra Direct, NSW, Australia. In Victoria, bird capture and banding was conducted over 20 days between winter 2008 and summer 2008/09. Colour-banding schema 07 (55) was used, adding a master colour-band to allow quick confirmation of each individual’s original capture site during field observations.

After banding, blood was collected from the right jugular vein and spotted onto Whatman filter paper (Sigma-Aldrich, N.S.W, Aust.). The sample was air-dried and placed in an individual envelope before being refrigerated at 4°C on site. Within 36 hours, the samples were transported to the Animal Genetics Laboratory (University of Queensland, St Lucia, Australia) for DNA sexing. Plumage indicators (53) were used to classify birds into two age groups: adult (>1 year) and juvenile. After sampling, birds were released adjacent to the study site.

### 2.4 Data collection

At each site, data were collected during two observation periods, each comprising three consecutive observation days. Observation periods were conducted in autumn (April) and spring (September) 2008 at site 1 and spring (September/October) 2008 and autumn (April/May) 2009 at site 2a. A single observation period was conducted at site 2b in autumn (April) 2009 to test the assumption that birds foraging there were also members of the population using the foraging resources at site 2a. A minimum of six days separated the observation periods from the last capture effort. Two additional observation periods were conducted at site 1 after the completion of the data collection for this study; the presence of a bird during these additional observations confirmed it was still alive and validated its inclusion in the dataset for site 1. Autumn and spring were selected for observational data collection to minimise environmental variables (these seasons are similar in temperature and photoperiod) and incorporate the expected seasonal variation in bird foraging effort associated with the non-breeding and breeding seasons. In addition, these seasons span the partial migration of the study species (44, 52, 53).

#### 2.4.1 Foraging observations

Each observation day ran from 08:00 to 16:00, with data collected every 10 minutes. Preliminary fieldwork conducted pre-dawn to dusk indicated that it was rare for study species to be observed outside these times. During each 10-minute interval (observation interval), the entire feeding area was scan sampled (56) by the same observer, who followed a circuit transect, maintaining a line of sight <5m from the birds under observation. In Victoria, a second observer was required to assist counting during the holiday period, due to the larger area to be covered and the number of people and birds present.

All colour-banded birds observed foraging during each observation interval—including those that flew in— were recorded in the field data sheets. Birds were considered to be foraging if they were ingesting, manipulating or reaching for food (54), or located at a point where seed was available The data recorded included the time, species and unique colour-band set. Data recording for each observation interval took between one and eight minutes; there were no occasions when there were so many birds that the recording of one interval interfered with the start of the next.

#### 2.4.2 Seed selection and intake

Independently of the observation periods, seed selection was determined by offering a handful of commercial seed mix (Golden Cob Cockatiel Mix [Mars Birdcare, Wacol, Qld]—large seeds: grey-striped sunflower, safflower, hulled oats; small seeds: white french, japanese and shirohie millet, canary, linseed, canola and milo) on a handheld tray for 5 minutes (*n*=8). In preparation, each handful of seed was manually divided into groups for each type of large seed and as a combined group for small seeds. The mass of each group was recorded in grams to two decimal places. After birds fed from the tray, whole seeds and husks were divided into the respective groups and weighed separately.

Seed intake was estimated by filming individual Crimson Rosellas (*n*=5) and Australian King-Parrots (*n*=5) for one 10-minute interval under normal foraging conditions. The recordings were reviewed, noting the feeding point/s (hand/ground) and the birds’ seed intake rate (rapid, casual, constant, slow). In addition, the time spent feeding (% of interval), the number of times a bird moved between feeding points and the number of times a bird picked up (prehended) a large or a small seed were counted.

#### 2.4.3 Incidental observations

Additional observations of factors that contributed to the foraging opportunity were recorded. These included details about behaviours of the study species, numbers and behaviours of other species using the provisioned seed, and factors such as disturbances. Ambient temperatures (°C) were recorded for comparison with long-range average temperatures for the same time of the year.

#### 2.4.4 Bird counts and the number of people feeding birds

The total number of each study species observed foraging at the site (that is banded and unbanded birds) was counted, on average, every 30 minutes throughout the day, but more frequently when bird numbers were observed to be increasing. On the first day of each observation period, the total number of people feeding the birds (participants) was counted and recorded every hour.

### 2.5 Data management

Field data were transferred to a Microsoft Excel spreadsheet. Individual birds, which were specific to a site, were included in the dataset if they met the following criteria: 1. The unique colour-band combination noted in the field data sheet cross-referenced to an individual in the capture and banding data, positively identifying the bird; 2. The bird was recorded foraging during any observation period after capture (including birds recorded foraging during additional post-study observations at site 1) and was therefore known to be alive. The data for three birds was discarded due to a band falling off (*n*=2) and a recording error in the field (*n*=1).

The following metrics (demonstrated in S1 Fig.) were used to summarise the observed foraging effort for each bird over the three days of an observation period: 1. Frequency-days (Fd) – the total number of days in which a bird was observed foraging during an observation period (0, 1, 2, or 3; those birds recording 0 were observed foraging during the previous, or a subsequent observation period). 2. Frequency-visits (Fv) – the number of visits a bird made each day during an observation period. A separate visit was counted if a bird was absent for ≥ 20 minutes before returning to feed. This criterion was included to prevent brief fly-offs resulting in an overestimation of the number of visits. 3. Visit duration (Vd) – because each bird’s visit duration was not timed, the value of Vd was assumed to be equal to 10 minutes times the number of observation intervals in which the bird was recorded foraging for that visit. If, during one visit, a bird was absent for one observation interval, that interval was not included in the calculation of Vd. 4. Daily duration (Dd) – the total number of foraging minutes recorded for a bird each day. The value of Dd was assumed to be equal to 10 minutes times the number of observation intervals in which the bird was observed foraging. 5. Foraging score (Fs) – calculated using the formula (Σ Dd for an observation period/10) x Fd. This metric was calculated to represent a bird’s foraging effort over an observation period, accounting for both its’ foraging frequency and duration.

### 2.6 Data analyses

#### 2.6.1 Data summaries

Site datasets for each observation period were separated into species and individual bird metrics were summarised for each species. To assist with visual comparison of the two sites’ data summaries, Victoria’s data is presented in the order of the seasons, not in the chronological order of collection. Given the total number of birds (n value) differed for each metric, all metrics were expressed as a relative frequency (percentage) to allow comparisons between species, seasons and sites. The relative frequencies of the possible outcomes for Fd and Dd were calculated including observed values of zero. The relative frequencies for Fv and Vd were calculated using the maximum value a bird recorded over the observation period. These calculations did not include observed values of zero. Foraging scores were summarised using measures of central tendency—including the mean, standard error and median. Each bird’s foraging effort was categorised into one of 6 Foraging classes (Fc): Fc0=Fs0, Fc1=Fs1-25, Fc2=Fs26-50, Fc3=Fs51-75, Fc4=Fs76-100, and Fc5=Fs100-125. The relative frequency (%) of the number of birds in each class were calculated for each species.

#### 2.6.2 Restricted Maximum Likelihood Modelling

The remaining statistical analyses of the birds’ foraging scores were conducted using GenStat for *Windows* 18^th^ Edition. (VSN International, Hemel Hempstead, UK) with significance set at a *p*-value of less than 0.05. Non-categorised foraging score data (response variable) as a continuous variable was analysed using a Restricted Maximum Likelihood (REML) Model. Fixed effects considered for inclusion were site, season, species, sex, age, holiday/not holiday and fly-off events (total), with random effects of bird identification and day. The data were square root transformed to meet the underlying assumption of normality for the REML Model. Predicted means and standard errors were obtained from the modelling and back-transformed onto the original scale.

Univariate analyses were conducted on each fixed effect to determine initial significance. Any term with a *p*-value less than 0.25 was considered for inclusion in the multivariate model, whereby a stepwise backwards elimination approach was used to determine a final model in which all fixed effects were significant. All two and three-way interactions were tested, and Least Significant Differences were obtained to explore pairwise differences.

#### 2.6.3 Estimated energy intake

The number of kilojoules (kJ) ingested by Crimson Rosellas and Australia King-Parrots that were monitored over a 10-minute interval was estimated by converting the number of prehended seeds into volume (ml), multiplied by the energy density (kJ/ml) for each seed type. If a bird consumed both large and small seeds, the energy contribution of each was calculated independently then summed. Energy densities were calculated based on average published values for large seeds (13.90kJ/ml) and small seeds (12.38kJ/ml) (57). The percentage of the daily energy requirements consumed by each bird was calculated by dividing the kilojoules by the species’ daily energy requirement. The daily energy requirement of wild Crimson Rosellas has been reported to be 157kJ (54). The daily energy requirement for wild Australian King-Parrots has not been reported; it was therefore estimated from the daily energy requirement for Crimson Rosellas and metabolic scaling (58). The daily energy requirements for an Australian King-Parrot at the lower end of their weight range (210g) was calculated to be 237kJ.

#### 2.6.4 Population estimates

Population estimates for birds using the feeding site were calculated for the study species using a methodology appropriate for the different sample sizes and extent of resampling data. Population estimates were calculated for Crimson Rosellas using the Jolly-Seber Method (59). The model was populated using resighting data for colour-banded birds and a single count of unbanded birds (total count minus the number of banded birds observed for that observation interval). The values were taken from three discrete time points per day for three days. The discrete time points fell within the following time periods: 08:00 to 10:30, 10:40 to 13:10 and 13:20 to 15:50. As data were derived from observations (not recaptures) no adjustment was made for capture bias. A population estimate was calculated for each of the nine time points; the average and the maximum of these were reported. Population estimates were calculated for Australian King-Parrots using the Peterson Method, applying the appropriate estimator from Bailey (59). The maximum counts of colour-banded AKP from an observation period were used to approximate the maximum population estimate with a 95% confidence interval.

## 3. Results

### 3.1 Numbers of colour-banded birds

#### 3.1.1 Queensland and Victoria (2a)

Over the course of the entire study, a total of 269 birds were colour-banded (197 Crimson Rosellas and 72 Australian King-Parrots). The number of birds that were colour-banded by the time of each observational period, the number of these that were sighted in each observation period or follow up observation periods, and the number of birds that were observed foraging each observation period are shown in Table 1. The demographics (age and sex) of the colour-banded birds known to still frequent the site were as follows: Crimson Rosellas—QLD adult male (am)=28, adult female (af)=18, adult unknown (au)=29, juvenile male (jm)=3, juvenile female (jf)=4, juvenile unknown (ju)=2; VIC am=19, af=14, jm=4, jf=3; Australian King-Parrots—QLD am=8, af=3, au=2, jm=0, jf=7, ju=3; VIC am=2, af=2, jm=1, jf=2. What happened to the banded birds that were never observed again is not known. They may have still been using the feeding sites at times other than the observation periods, or they may have died or emigrated.

**Table 1:** Listed are the total number of Crimson Rosellas and Australian King-Parrots that had been colour-banded by the time of the autumn and spring observation periods at wild bird feeding sites in Queensland and Victoria, showing the number of birds in each dataset and the number observed foraging during each three-day observation period. To assist comparison, the numbers for Victoria are not shown in the chronological order of collection.

|  | Queensland |  |  |  | Victoria |  |  |  |
| --- | --- | --- | --- | --- | --- | --- | --- | --- |
|  | Crimson Rosellas |  | Aust. King-Parrots |  | Crimson Rosellas |  | Aust. King-Parrots |  |
|  | Autumn<br>2008 | Spring<br>2008 | Autumn<br>2008 | Spring<br>2008 | Autumn<br>2009 | Spring<br>2008 | Autumn<br>2009 | Spring<br>2008 |
| Total number of colour-banded birds at the time of an observation session | 80 | 99 | 35 | 46 | 98 | 66 | 26 | 17 |
| Total number of colour-banded birds known to still frequent the site | 67 | 84 | 20 | 23 | 40 | 31 | 7 | 6 |
| Number of colour-banded birds observed foraging during the observation session | 58 | 79 | 12 | 17 | 29 | 26 | 2 | 5 |

#### 3.1.2 Birds’ movement between sites 2a and 2b, in Victoria

Observations conducted at site 2b confirmed that Crimson Rosellas moved between feeding sites 2a and 2b to use provisioned seed. By autumn, 65 Crimson Rosellas were colour-banded at site 2a and 33 at site 2b in Victoria. Nine Crimson Rosellas observed foraging at site 2b in autumn 2009 were originally caught at site 2a and one Crimson Rosella caught at 2b was observed foraging at site 2a. Twenty-two Australian King-Parrots were colour-banded at site 2a and four at site 2b. This species was not observed foraging at the alternate sites during the observation periods.

### 3.2 Foraging metrics

#### 3.2.1 Frequency-days

The percentage of birds observed foraging for 0, 1, 2, or 3 days is shown in Fig.1, illustrating the differences between species and seasons for each site. Not all of the birds were observed using the foraging resource every day, and some did not during the observation period. In Queensland, where Crimson Rosellas were observed using the feeding site often, only 49.25% and 51.19% were observed foraging every day in autumn and spring, respectively. In total (over both sites and seasons), 13.51% of Crimson Rosellas used the foraging resources zero days, 19.37% 1 day, 20.27% 2 days and 46.85% 3 days. Of the Australian King-Parrots, 35.71% used the foraging resources zero days, 17.86% 1 day, 19.64% 2 days and 26.79% daily (data not shown).

#### 3.2.2 Frequency-visits

The percentage of birds that fed at a site a maximum of one to eight times a day during each observation period is shown in Fig. 2. Of the birds observed using the foraging resource, the majority of Crimson Rosellas were recorded making between one to three visits in a day, while the majority of Australian King-Parrots made only made one or two visits in a day. The highest number of visits recorded for a bird was eight, for one Crimson Rosella in Queensland.

**Figure 1:**
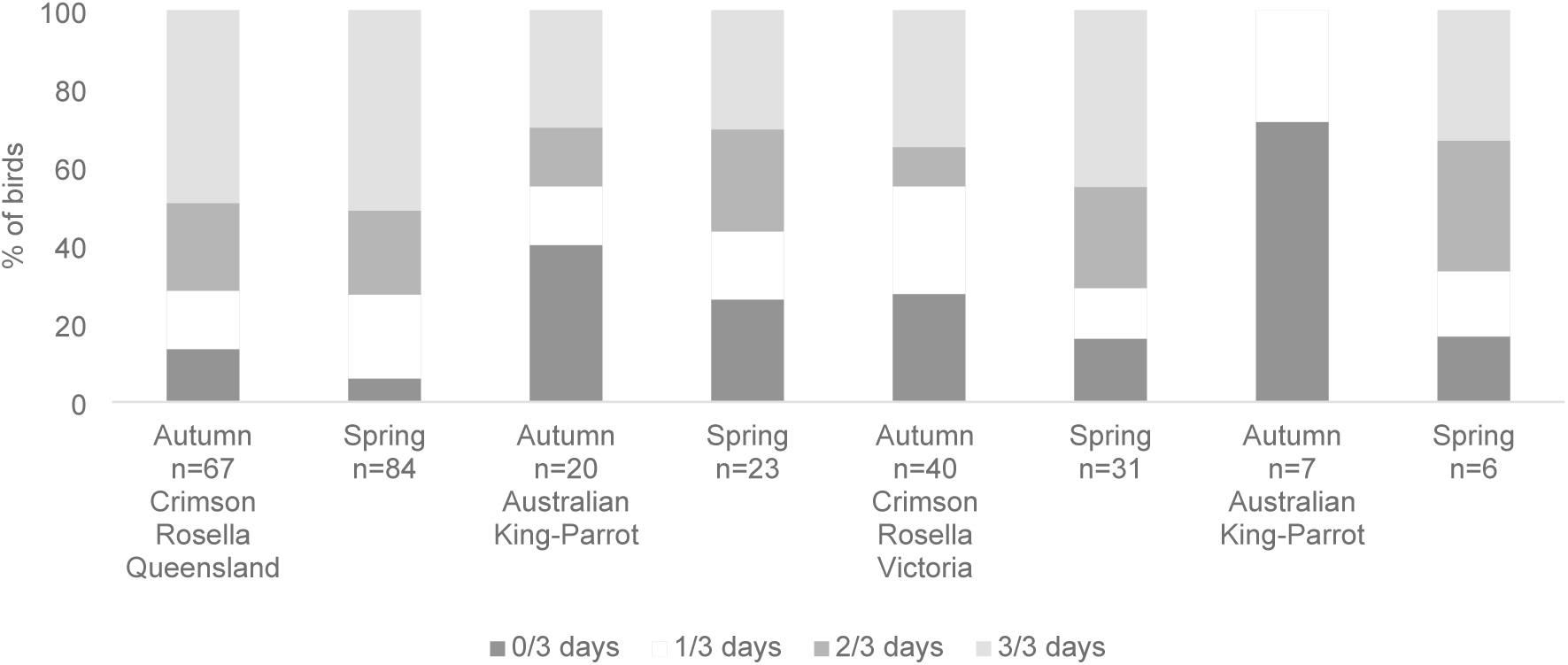
The percentage of colour-banded Crimson Rosellas and Australian King-Parrots that fed at a Queensland and Victorian feeding site on zero, one, two or three days during a three-day observation period in spring and autumn.

**Figure 2:**
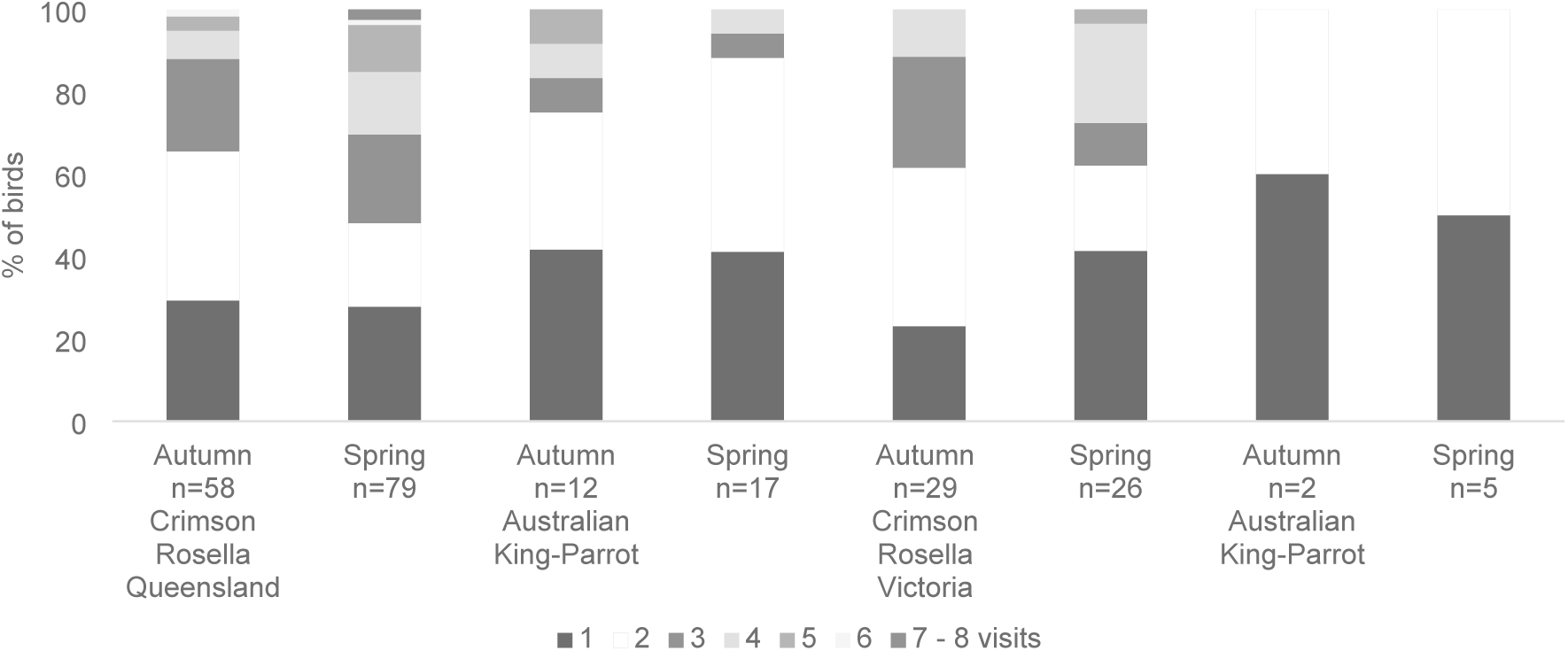
The percentage of colour-banded Crimson Rosellas and Australian King-Parrots that fed at a Queensland and Victorian feeding site a maximum of one to eight times a day during a three-day observation period in spring and autumn. Given the small percentage of birds recorded making a maximum of 7 and 8 visits in a day, these outcomes were pooled.

#### 3.2.3 Visit duration

The percentage of birds that fed at a site for a maximum visit duration of 10 to 120 minutes during each three-day observation period is shown in Fig. 3. Overall, most individual visits lasted 20 minutes or less. Only a small percentage of visits lasted for longer than 30 minutes; Australian King-Parrots in Queensland during autumn were an exception. The longest visit duration recorded for a Crimson Rosella was 120 minutes in Queensland and 70 minutes in Victoria. The longest visit duration recorded for Australian King-Parrots was 60 minutes in Queensland and 20 minutes in Victoria.

**Figure 3:**
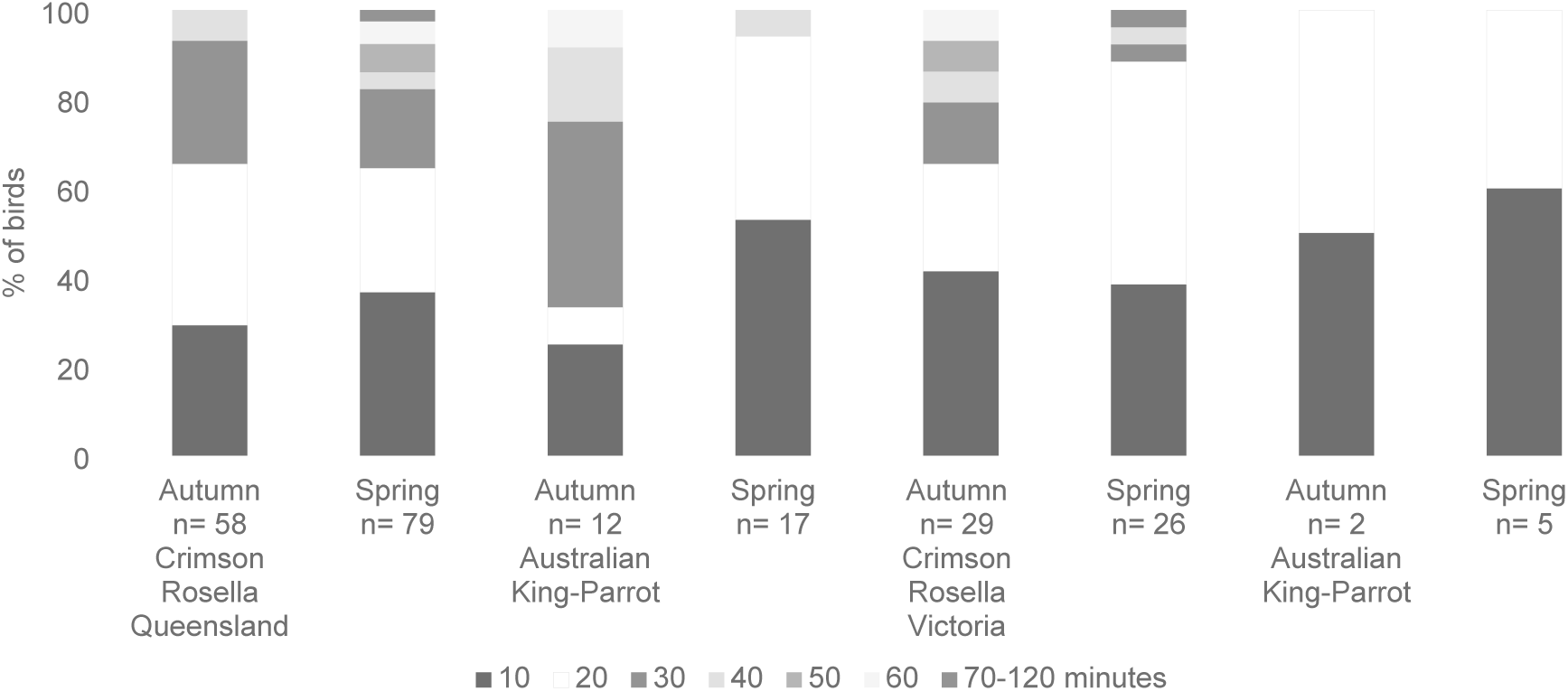
The percentage of colour-banded Crimson Rosellas and Australian King-Parrots that fed at a Queensland and Victorian feeding site for a maximum visit duration of 10 to 120 minutes during a three-day observation period in spring and autumn. Given the small percentage of birds that recorded a maximum visit duration of 70 to 120 minutes, these outcomes were pooled.

#### 3.2.4 Daily duration

The percentage of birds that recorded a daily duration between 0 and 160 minutes over each three-day observation period is shown in Fig. 4. As a result of the birds that were not observed using the site on one or more days during an observation period, the most frequently recorded daily foraging duration was zero. Australian King-Parrots in Victoria during spring were an exception. Across both observation periods, over 75% of birds were recorded using the foraging resources for 30 minutes or less each day [75.96% CR (*n*=666) and 81.55% AKP (*n*=168)]. The longest daily duration recorded for Crimson Rosellas was 160 minutes at both locations. Longer daily durations were recorded by Crimson Rosellas during spring in Queensland and during autumn in Victoria. The longest daily duration recorded for Australian King-Parrots was 150 minutes during autumn in Queensland and 30 minutes during spring in Victoria.

**Figure 4:**
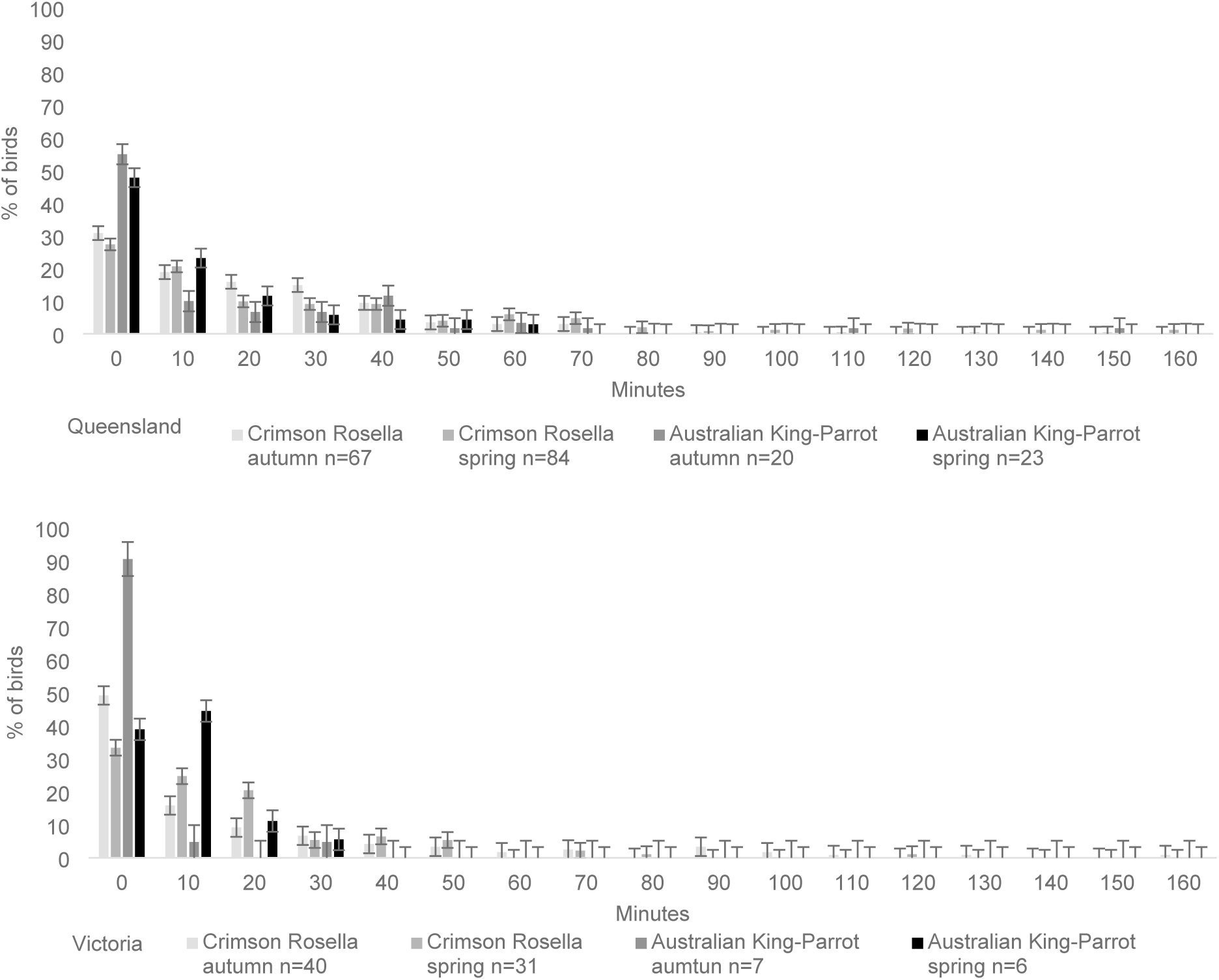
The percentage of colour-banded Crimson Rosellas and Australian King-Parrots that recorded a daily duration between 0 and 160 minutes at a Queensland and Victorian feeding site over a three-day observation period in spring and autumn. Error bars represent one standard error.

#### 3.2.5 Foraging scores

A summary of the birds’ foraging scores and the percentage of birds in each class are provided in Table 2. Overall, birds’ foraging scores ranged from 0 to 117. The percentage of birds in each Foraging class was as follows: Fc0=18.3%, Fc1=55.4%, Fc2=19.8%, Fc3=4.3%, Fc4=1.1%, Fc5=1.1%. The foraging effort of 5 birds—one from each class—is shown in S2 Fig. to demonstrate the relationship between foraging effort, foraging score and class.

**Table 2:** A summary of birds’ foraging scores (Fs) and the percentage of birds with foraging scores in each class, calculated for colour-banded Crimson Rosellas and Australian King-Parrots that fed at a Queensland and Victorian feeding site during a three-day observation period in spring and autumn. Foraging class (Fc) 0=Fs 0, Fc1=Fs1-25, Fc2=Fs 26-50, Fc3=Fs51-75, Fc4=Fs76-100, Fc5=Fs101-125.

|  | Queensland |  |  |  | Victoria |  |  |  |
| --- | --- | --- | --- | --- | --- | --- | --- | --- |
|  | Crimson Rosella |  | Aust. King-Parrot |  | Crimson Rosella |  | Aust. King-Parrot |  |
|  | Autumn | Spring | Autumn | Spring | Autumn | Spring | Autumn | Spring |
| *n* | 67 | 84 | 20 | 23 | 40 | 31 | 7 | 6 |
| **Foraging scores** |  |  |  |  |  |  |  |  |
| Mean | 18.26 | 26.06 | 24.17 | 12.76 | 22.79 | 15.74 | 2 | 7.4 |
| Standard error | 1.79 | 2.86 | 7.49 | 2.83 | 5.06 | 2.95 | 1 | 2.38 |
| Median | 17 | 18 | 18 | 10 | 8 | 12 | 2 | 8 |
| **Foraging class (%)** |  |  |  |  |  |  |  |  |
| **0** | 13.43 | 5.95 | 40.00 | 26.10 | 30.00 | 16.20 | 71.50 | 16.70 |
| **1** | 64.18 | 51.19 | 35.00 | 65.20 | 42.50 | 74.20 | 28.60 | 83.40 |
| **2** | 20.90 | 30.95 | 20.00 | 8.80 | 15.00 | 9.60 | 0 | 0 |
| **3** | 1.49 | 7.14 | 0 | 0 | 10.00 | 0 | 0 | 0 |
| **4** | 0 | 2.38 | 5.00 | 0 | 0 | 0 | 0 | 0 |
| **5** | 0 | 2.38 | 0 | 0 | 2.50 | 0 | 0 | 0 |

Univariate analyses indicated that the predicted mean foraging score for the pooled sample of scores was significantly higher in Queensland than in Victoria, higher in spring than in autumn, higher for Crimson Rosellas than Australian King-Parrots and higher for adults than juveniles, as listed in Table 3. There was no significant difference for gender, holiday/not holiday, or fly-off events.

**Table 3:** Univariate analysis to identify predictor variables influencing the foraging score.

| Variable | Category | Predicted mean | Standard error | *p* value |
| --- | --- | --- | --- | --- |
| Site | Queensland | 35.69 | 1.21 | **0.011** |
|  | Victoria | 14.64 | 1.34 |  |
| Season | Spring | 34.95 | 1.19 | **<0.001** |
|  | Autumn | 20.57 | 1.19 |  |
| Holiday / not holiday | Holiday | 24.75 | 1.20 | 0.215 |
|  | Not holiday | 29.11 | 1.19 |  |
| Species | Crimson Rosella | 35.91 | 1.19 | **<0.001** |
|  | Australian King-Parrot | 8.71 | 1.42 |  |
| Sex | Male | 27.06 | 1.29 | 0.628 |
|  | Female | 22.51 | 1.33 |  |
| Age | Adult | 33.89 | 1.19 | **0.005** |
|  | Juvenile | 10.52 | 1.44 |  |
| Fly-off events (total) | within 10 minutes | 29.49 | 1.19 | 0.172 |
|  | within 25 minutes | 25.89 | 1.19 |  |
|  | within 40 minutes | 22.13 | 1.25 |  |

The final multivariate model included the fixed effects of site (p=0.008), season (p=<0.001) and species (p=<0.001). Age (p=0.083), holiday/not holiday (p=0.74) and fly-off events (p=0.172) were not significant in the multivariate model. A significant three-way interaction existed between species, season and site (p=<0.001), as demonstrated in Fig. 5.

**Figure 5:**
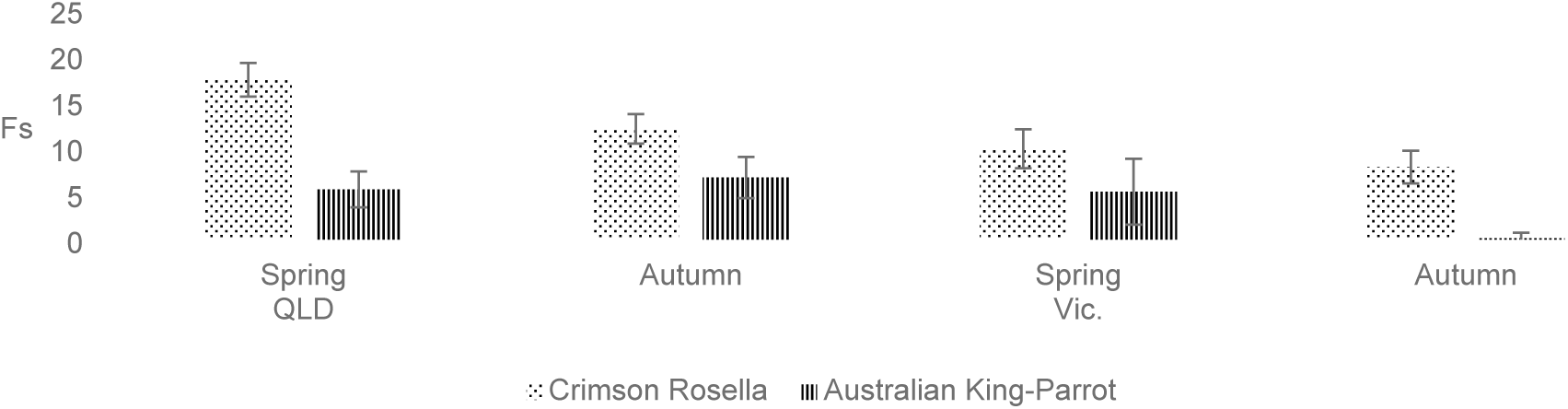
Three-way interaction between species, season and site affecting foraging scores.

Further investigation into the significant (p,0.001) 3-way interaction revealed the following differences. There was no significant variation in Australian King-Parrots’ foraging scores in spring across sites (Qld 5.5±1.9, Vic. 5.2±3.6). Crimson Rosellas’ foraging scores in spring were significantly higher in Queensland as compared to Victoria (17.4±1.8 vs 9.9±2.1). In autumn, Australian King-Parrots had significantly higher foraging scores in Queensland than Victoria (6.8±2.2 vs 0.15±0.6). In autumn, there was no significant variation in foraging scores for Crimson Rosellas across the sites (Qld 12.05±1.6 vs Vic. 7.9±1.8).

In Queensland, Crimson Rosellas had significantly higher foraging scores in spring as compared to autumn (17.4±1.8 vs 12.05±1.6). There was no variation in the Australian King-Parrots foraging scores in Queensland between seasons (5.5±1.9 vs 6.8±2.2). In Victoria, Australian King-Parrots had significantly higher foraging scores in spring as compared to autumn (5.2±3.6 vs 0.15±0.6) and Crimson Rosellas showed no difference in foraging scores between seasons (9.8±2.1 vs 7.9±1.8). While statistically significant differences were observed within the Australian King-Parrots results, the high standard errors, which reflect the low sample size, indicate these findings should be interpreted with caution.

### 3.3 Seed selection and percentage of daily energy requirements consumed

#### 3.3.1 Seed selection

Preferential seed selection was observed during each of the 5-minute intervals that birds were offered the seed mix. Birds favoured sunflower and safflower seeds, followed by hulled oats. Preferential seed selection was consistent for Australian King-Parrots, but Crimson Rosellas would intermittently prehend smaller seeds. In each interval, all of the sunflower seeds were consumed and in six intervals all of the safflower seeds were consumed. In the two intervals that safflower seeds were not depleted, fewer than 10% remained. In each interval between 50-80% of the oats were consumed. Small seeds were always consumed, but at less than 10% of the mass offered. Within the 5-minute intervals, between 1-11 birds would feed from the tray. Competition was observed between birds of the same species and between Crimson Rosellas and Australian King-Parrots; this regularly resulted in feeding being disrupted and birds being displaced from the tray.

Two occasions when visitors brought their own supply of sunflower seeds provided the opportunity to observe birds feeding on this seed type. On one occasion, a person hand fed an Australian King-Parrot for five minutes. A feed intake rate of 1 prehended seed per 3 seconds was calculated, resulting in prehension of approximately 200 seeds. On another occasion, a person was observed hand feeding black sunflower seeds to two Australian King-Parrots. One bird fed for one minute and the other fed for fifty seconds before they were displaced. The first bird consumed 28 seeds (one seed/2.1secs) and the second bird consumed 14 seeds (one seed/3.5secs).

#### 3.3.2 Estimates for the percentage of daily energy intake

A summary of data for individual birds’ foraging for a 10-minute interval is presented in Table 4. In a 10-minute interval, Crimson Rosellas and Australian King-Parrots consumed between 1.73-62.91% and 6.84-88.54% of their daily energy requirements, respectively. Fully distended crops were regularly noted for both Crimson Rosellas and Australian King-Parrots during blood collection. Depending on which seeds were consumed, a full crop for either species would represent between 79% (small seeds) and 88% (large seeds) of their daily energy requirements.

**Table 4:** A summary of individual birds’ foraging data for a 10-minute interval at a feeding site in Queensland. Crimson Rosellas (*n*=5) and Australian King-Parrots (*n*=6)

| Bird # | Time spent feeding | Feeding point | # of times bird moved | Feed intake rate | Small seeds | mls | kJ | Large seeds | mls | kJ | Total kJ | Percentage of daily energy requirements consumed |
| --- | --- | --- | --- | --- | --- | --- | --- | --- | --- | --- | --- | --- |
| **Crimson Rosellas** |  |  |  |  |  |  |  |  |  |  |  |  |
| 1 | 49% | ground | 1 | casual | 242<sup>3</sup> | 2.81 | 34.84 | 0 | 0 | 0 | 34.84 | 22.19% |
| 2 | 100% | ground | 0 | constant | 647<sup>3</sup> | 7.52 | 93.14 | 0 | 0 | 0 | 93.14 | 59.32% |
| 3 | 15% | hand | 2 | casual | 0 | 0 | 0 | 18 | 0.9 | 12.51 | 12.51 | 7.97% |
| 4 | 9.2% | both | 1 | rapid | 14 | 0.16 | 2.02 | 1 | 0.05 | 0.695 | 2.71 | 1.73% |
| 5 | 31.2% | ground | 1 | rapid | 15 | 0.17 | 2.16 | 139<sup>1</sup> | 6.95 | 96.61 | 98.76 | 62.91% |
| **Australian King-Parrots** |  |  |  |  |  |  |  |  |  |  |  |  |
| 1 | 75% | both | 3 | rapid | 337<sup>3</sup> | 3.92 | 48.51 | 37 | 1.85 | 25.72 | 74.23 | 31.32% |
| 2 | 29.2% | hand | 5 | slow | 0 | 0 | 0 | 30 | 1.5 | 20.85 | 20.85 | 8.8% |
| 3 | 67.5 | hand | 11 | rapid | 76 | 0.88 | 10.94 | 156 | 7.8 | 108.42 | 119.36 | 50.36% |
| 4 | 64% | hand | 2 | rapid | 33 | 0.38 | 4.75 | 130 | 6.5 | 90.35 | 95.10 | 40.13% |
| 5 | 20.2% | ground | 7 | rapid | 74 | 0.86 | 10.65 | 8 | 0.4 | 5.56 | 16.21 | 6.84% |
| 6<sup>2</sup> | 50% | hand | 1 | rapid | 0 | 0 | 0 | 200<sup>3</sup> | 10 | 139 | 139 | 88.54% |

<sup>1</sup> Bird being fed at the ground, by a child using their own seed mix.
<sup>2</sup> An additional Australian King-Parrot that was being hand fed black sunflower seeds, was opportunistically added to this data set.
<sup>3</sup> Calculations represent maximum feed intake, based on whole seeds.

### 3.4 Summary of incidental observations

#### 3.4.1 Factors observed to influence the foraging opportunity

The foraging opportunity was influenced by the type and quantity of whole seeds available at each feeding point (hand/ground). As seeds were hulled, there was an increasing ratio of husks to whole seeds. A bird’s feeding efficiency would be affected by the number of times it prehended husks. The ratio of whole seeds to husks at the hand would remain reasonably high as people continually replenished the supply. An examination of waste (seeds and husks collecting on the ground) from a 10cm square revealed more than half of the material was husks and small seeds represented the bulk (>95%) of the whole seeds, but these quantities would constantly be changing. Both of the study species fed from peoples’ hands and on the ground. Taking into consideration all of the foraging observations of Crimson Rosellas (*n*=1671) and Australian King-Parrots (*n*=235), Crimson Rosellas and Australian King-Parrots were observed feeding at the hand 8.98% and 85.96% of the time, respectively.

Observations indicated that feeding efficiency was also influenced by a bird’s feed intake rate and level of vigilance, the amount of time a bird spent walking or flying between feeding opportunities, the level of disturbance, the time a bird spent taking shelter in response to a threat (often preceded by alarm calls triggered by loud noises or predatory birds), the number of birds feeding and the level of competition.

Seed availability and competition was also influenced by other species of birds that used the provisioned seed. In Queensland, the primary incidental species were Australian Brush-turkey (*Alectura lathami)* (max. autumn 25, spring 13) and Red-browed Finch (*Neochmia temporalis)* (autumn/spring max.12). Both species fed throughout the day on seed that collected at the ground and few incidents of interspecies competition were observed. Australian Brush-turkey numbers were generally low (≤5) but increased in the afternoon when people and the majority of study species had departed. In Victoria, other species observed foraging at the site were Sulphur-crested Cockatoo (*Cacatua galerita)* (max. autumn 128, spring 195), Galah (*Eolophus roseicapilla)* (max. autumn 30, spring 35) and Long-billed Corella (*Cacatua tenuirostris)* (max. autumn 0, spring 4). All of these species fed at the ground and Cockatoos and Galahs regularly fed directly from peoples’ hands. Cockatoos regularly outnumbered other species present at this site and were commonly observed displacing all other species.

#### 3.4.2 Climatic conditions

The temperatures at both sites were within the normal temperature range for the time of year, with spring at site 2 being a slight exception—the average daily maximum temperature was 3.5°C higher than the long-term average for this month. In Queensland, the average daily maximum temperature was 21°C for both seasons (long-term averages April 22°C, September 21°C (60)). In Victoria, the average daily maximum temperatures for autumn and spring were 13.4°C and 20.5°C, respectively (long-term averages May 14°C, October 17°C (61)).

### 3.5 Numbers of birds and participants

#### 3.5.1 Counts

Counts of the birds using the foraging resource and the people feeding them fluctuated throughout the day at both sites, as shown in S3 Fig.. Participant numbers were also observed to increase on weekends (data not shown) and during holiday periods. At both sites, people were generally present in larger numbers from mid-morning when tourist buses arrived. Sometimes there were more people than birds present, and occasionally, there were small numbers of participants feeding large numbers of birds. There were intervals when one or both species were absent. Crimson Rosellas were consistently present in greater numbers than Australian King-Parrots.

#### 3.5.2 Maximum counts and population estimates

From the numbers of studied species counted using the feeding site (banded and unbanded birds), the maximum recorded each observation period is shown in Table 5, along with the population estimates for each species. Population estimates were on average 4.05 and 3.16 times greater than the maximum counts for Crimson Rosellas and Australian King-Parrots (Qld only), respectively. Both the maximum counts and population estimates for Crimson Rosellas increased in spring. Maximum counts of Australian King-Parrots using the foraging resource were relatively constant, but a seasonal increase was observed in the population estimate for the Queensland site in spring. Due to the small sample size and low number of foraging observations recorded for Australian King-Parrots in Victoria during autumn, it was not possible to produce a population estimate.

**Table 5:** The maximum numbers and population estimates of Crimson Rosellas and Australian King-Parrots recorded foraging at a Queensland and Victorian wild bird feeding site during a three-day observation period in autumn and spring. CI = Confidence interval.

|  | Crimson Rosellas |  | Australian King-Parrots |  |
| --- | --- | --- | --- | --- |
|  | Max. # | Population estimate<br>Average (max.) | Max. # | Population estimate<br>Max. (95% CI) |
| **Queensland** |  |  |  |  |
| Autumn | 70 | 289 (795) | 10 | 29 (26, 32) |
| Spring | 130 | 577 (1968) | 11 | 45 (42, 40) |
| **Victoria** |  |  |  |  |
| Autumn | 80 | 235 (940) | 10 | - |
| Spring | 105 | 492 (2357) | 10 | 25 (22, 29) |

## 4. Discussion

This study is the first to examine how the provisioned seed offered at wild bird feeding sites influences the behaviour of two psittacine species in Australia. At both sites, the availability of provisioned seed resulted in a marked increase in the number of Crimson Rosellas foraging together at a single location, as compared to what would occur in the wild. Under natural conditions, Crimson Rosellas typically forage in small flocks ranging from two to five birds and would not consistently forage with Australian King-Parrots (52).

Additionally, once family groups separate, adult and juvenile Crimson Rosellas would typically segregate and forage in different habitats (62). At the feeding sites, up to 130 (spring) and 70 (autumn) Crimson Rosellas were observed foraging at one time at the Queensland site; a similar result was found at the Victorian site with maximums of 105 in spring and 80 in autumn. Additionally, adult and juvenile Crimson Rosellas were seen to freely intermingle and both Crimson Rosellas and Australian King-Parrots were regularly present at the feeding sites at the same time, along with various other species that would not forage together under natural conditions. The extent of altered foraging behaviour at the feeding sites can be further appreciated given that the estimated number of Crimson Rosellas cycling through them was 2.9 to 4.7 times higher than the maximum number of birds seen at any one time during each observation period. A conservative estimate of the numbers of Crimson Rosellas foraging at the Queensland and Victorian sites at peak use (in spring) was 577 and 492, respectively.

To understand the potential impacts of birds being offered provisioned seed at wild bird feeding sites, we sought to verify if birds foraged at the sites every day, to determine how often and how long they foraged on visit days, and to estimate the proportion of birds’ daily energy requirements being met by the seed consumed. If birds met most or all of their energy requirements by foraging at wild bird feeding sites, they would be more likely to be negatively impacted by sudden changes in seed availability. It has also been postulated that a constant source of provisioned feed could alter birds natural movement patterns and result in juveniles not learning natural foraging skills (16, 20).

In this study, a spectrum of use was observed in the Crimson Rosellas. Categories of use could be broken down into birds that did not forage at a feeding site during a three-day observation period—but were seen using it at other times, birds that foraged at a feeding site one or two days out of three, and birds that foraged at a feeding site every day. In Queensland over both seasons, the number of birds that did not use the feeding site during the observation periods was small: <15%. Thirty to 45% used the feeding site one or two days out of three and approximately half used the feeding site every day. Given that the feeding site in Queensland was surrounded by wilderness, on days that Crimson Rosellas did not use the feeding site they would have been obtaining their diet from natural food sources. In Victoria, the percentage of Crimson Rosellas that did not use the feeding site or used it only one or two days was higher. These birds could potentially have been using the time away from the feeding site to feed on natural food sources; however, they could have fed at a nearby feeding site or backyard feeders—that were not monitored.

On days that Crimson Rosellas used the provisioned seed, the number of times a day they visited also varied between individuals. Just under 30% of the Queensland Crimson Rosellas in both seasons only visited the feeding site once a day, between 20% (spring) and 40% (autumn) visited the feeding site twice a day and 35% (autumn) and 55% (spring) visited the feeding site three of more times a day. Overall, frequency of visits to the feeding site in Victoria was comparatively lower in both seasons; this finding is consistent with the birds’ opportunity to make use of other feeding sites. The visit duration of the majority of Crimson Rosellas using a feeding site in both seasons was 10 to 30 minutes. Maximum daily durations at the feeding sites were commonly 50 minutes or less. In one ten-minute observation interval, it was possible for a Crimson Rosella to consume between 2-63% of their daily energy requirements.

Birds that consumed small quantities of seed, visiting for short periods of time either once or twice a day were more likely to be maintaining their use of natural foraging resources and using the feeding site as one of a range of resource patches. When foraging naturally, Crimson Rosellas regularly move from one foraging opportunity to another (52). This natural behaviour may influence seed intake at a feeding site, with birds consuming quantities in proportion to the fullness of their crop when they arrive at the site. It is likely that Crimson Rosellas consuming larger quantities of seed and either partially or completely filling their crop (gaining approximately 79-88% of their daily energy requirements) during one or more visits could readily be meeting a large portion, or even all of their energy needs by consuming the provisioned seed. In this situation, higher levels of feed intake would reduce the birds’ dietary diversity, although the provisioned seed is unlikely to be used as the sole food resource. In a study of the feeding ecology of a closely related species, the Adelaide Rosella (*Platycercus elegans adelaidae*), the crop contents revealed that no matter how abundant a food source was birds did not consume it exclusively, with evidence of four to eight different foods in sampled crops (63). When foraging naturally, Crimson Rosellas normally spend between 37-46% of the day foraging (52). For birds gaining the majority of their daily energy requirements at a wild bird feeding site, there would likely be a flow on effect on the birds’ time-activity budget, leaving more time available for other behaviours such as resting.

The population level analysis indicated that Crimson Rosellas’ foraging effort was comparable between sites and seasons, with spring in Queensland being the exception. In spring, there was a trend for more visits of longer duration in Queensland. The increased percentage of birds visiting feeding sites three or more times in this season and the increased foraging effort, may reflect the increased energy demands of breeding as some of the birds could be provisioning a mate or chicks. It is also likely that the increased number of birds using the site in spring would have increased competition and reduced birds’ feeding efficiency. This would require birds to increase their foraging effort to maintain the proportion of their energy requirements being obtained from the provisioned seed. In Victoria, a proportion of Crimson Rosellas increased their foraging effort in autumn, when daily temperatures were low. Some reports indicate this species increases its’ foraging effort during colder weather (52), but the opposite has also been reported (54).

The observations of Australian King-Parrots indicated that fewer numbers of this species utilised the feeding sites than Crimson Rosellas, with estimates of Australian King-Parrot numbers being 8-10% of the Crimson Rosella numbers in Queensland and 5% of the Crimson Rosella numbers in Victoria. The maximum number of Australian King-Parrots at the feeding sites (10 and 11) was also low compared to Crimson Rosellas. Species differences in the exploitation of provisioned food sources have previously been reported (1, 8, 64). A range of factors have been found to contribute to this variation, including the influence of the most abundant species using the feeder (65), behavioural tolerances (23), seed preferences and how the feed is presented (32, 66). Australian King-Parrots have been described as wary and while these parrots are known to forage on the ground they are more commonly observed foraging in trees (52). At the feeding sites, Australian King-Parrots were found to have a preference to feed directly from peoples’ hands, where competition was higher.

In addition to the discovery that fewer Australian King-Parrots used the feeding sites, they were also generally less likely to visit the feeding sites as compared to the Crimson Rosellas, and when they did, they visited fewer numbers of days, and averaged fewer visits per day. Despite this, Australian King-Parrots potentially gained a similar percentage of their daily energy requirements from the provisioned seed. Feeding from a persons’ hand, provided the advantage of preferential seed selection and the availability of a higher proportion of whole seeds. The observations of Australian King-Parrots’ seed intake confirmed this species could gain a half to full crop of seed in a ten-minute interval, providing between 50 and 89% of their daily energy requirements. In Victoria, Australian King-Parrots had to compete with large numbers of Sulphur-crested Cockatoo to feed at the hand, which may explain the low numbers of Australian King-Parrots at this location. Overall, the foraging conditions at both locations were likely more suited to Crimson Rosellas. Our data offer empirical evidence that concerns over wild bird feeding providing an advantage to one species over another (16) have merit. While Sulphur-crested Cockatoo was not a study species, incidental observations indicated that the wild bird feeding site in Victoria was likely to be providing an advantage to this species as well.

This study provides evidence that a proportion of the birds using the provisioned seed at both sites were likely to be dependent on the food source. A small percentage of birds (6.5% total, in classes 3-5—QLD CR aut.=1.49%, spr.=11.9%, AKP aut.=5%; Vic. CR aut.=12.5%) were recorded using the feeding sites heavily. All of these birds were observed foraging at a feeding site daily, recording visit durations up to 120 minutes and daily foraging durations up to 160 minutes. These birds may have had an increased energetic need as might be expected if breeding (67), unwell (1), travelling long distances (67), maintaining a higher body mass (64), or the site may have been central to the birds’ territory resulting in heavier usage (34). Irrespective of the reason for the increased foraging effort, it is highly likely that the birds in the higher foraging classes would be at risk, if the seed supply were suddenly reduced.

Authors have argued that wild bird feeding can disrupt the seasonal migration/movement patterns of birds (4, 16, 22). Our data does not support this hypothesis. At both sites, the bird counts and population estimates for Crimson Rosellas and Australian King-Parrots (excluding Australian King-Parrots at site 2) approximately doubled in spring compared to autumn, suggesting natural dispersal of subadults and non-resident adults of both species in the autumn was occurring. However, given that no base-line data is known for these sites, it is possible that a greater percentage of birds would have dispersed had the provisioned seed not been available.

Lastly, concerns that feeding could interfere with the development of natural food gathering behaviour (16) were not validated in this study. In studies of passerines at feeders, there have been mixed results for variation in foraging effort between age classes, with either no variation (31) or juveniles having higher foraging effort than adults (30). In our study, adults were found to have higher foraging effort at the study sites than juveniles. Therefore, we found no evidence to indicate that juvenile Crimson Rosellas or Australian King-Parrots were not developing the necessary skills for foraging on naturally occurring resources.

## Ethics and approvals

The University of Queensland Animal Ethics Committee approvals SVS/347/07/VAR/SF and SVS/671/08/VAR/SF covered bird capture and data collection in Queensland and Victoria. In Queensland, the Environmental Protection Agency (#WISP04635807) and O’Reilly’s Rainforest Retreat provided permission to conduct the study at site 1. Parks Victoria, the Department of Sustainability and Environment (#10004543) and the operator of the activity provided approvals for sites 2a and 2b in Victoria. The Australian Bird and Bat Banding Scheme approved the principle investigator’s R Class banding authority 2802 to conduct banding activities under project authority number 1379. The University of Queensland granted Human Ethics Committee approval and approved the fieldwork risk assessments.

## Acknowledgements

The authors extend their gratitude to the reviewers, Prof Peter O’Donoghue, Dr Robert Donnelly, A/Prof Lucio Fillipich, Queensland Parks and Wildlife Service, Grants on Sherbrooke, Healesville Sanctuary, Department of Environment and Conservation New South Wales, Tweed Bird Observers specifically Laurel Allsopp, Kathy Wilk, Steve McBride and Faye Hill, Jan and Bill Incol, Rita and Antonio Esposito, Dr Peter Holz, Dr Jenny McLelland, Dr Pam Whitely, Dr Anne Fowler and Dr Rosemary Booth, for providing extensive support to this project.

## Funding

This work was supported by O’Reilly’s Rainforest Retreat, Parks Victoria, Birds Queensland, Birds Australia, Bird Observers and Conservation Australia, Australian Wildlife Health Network, Annandale Bequest, Beaudesert Shire Council and Golden Cob.

## Conflict of interest

After this study, Michelle Plant assisted the management at both sites to implement feeding practices to meet regulatory agencies’ approval requirements and developed/sold equipment in support of this purpose.

## Data availability statement

Data for this article are available at <u>doi:10.5063/F1445JW7</u> (68)

## Supporting information

**Supplementary Figure 1:**
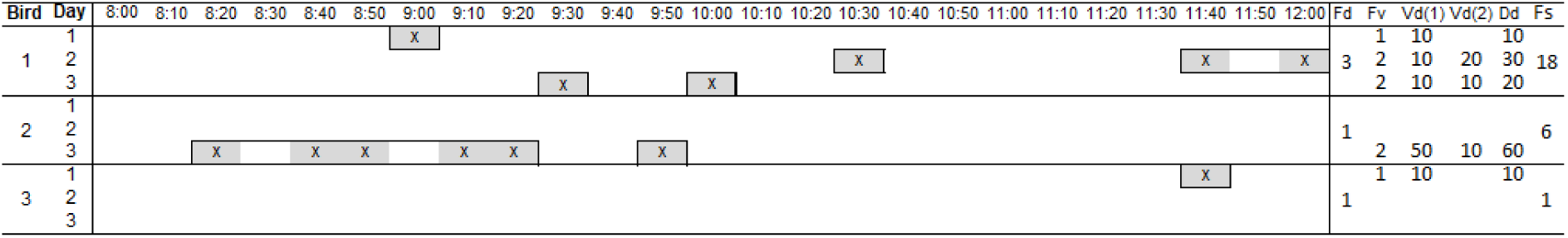
Graphic representation of how the foraging metrics (Frequency-days, Frequency-visits, Visit duration, Daily duration, and Foraging score) were obtained. For presentation purposes, the metrics shown only cover data from 08:00am to 12:00noon. The intervals considered to be a single visit are shown in boxes.

**Supplementary Figure 2:**
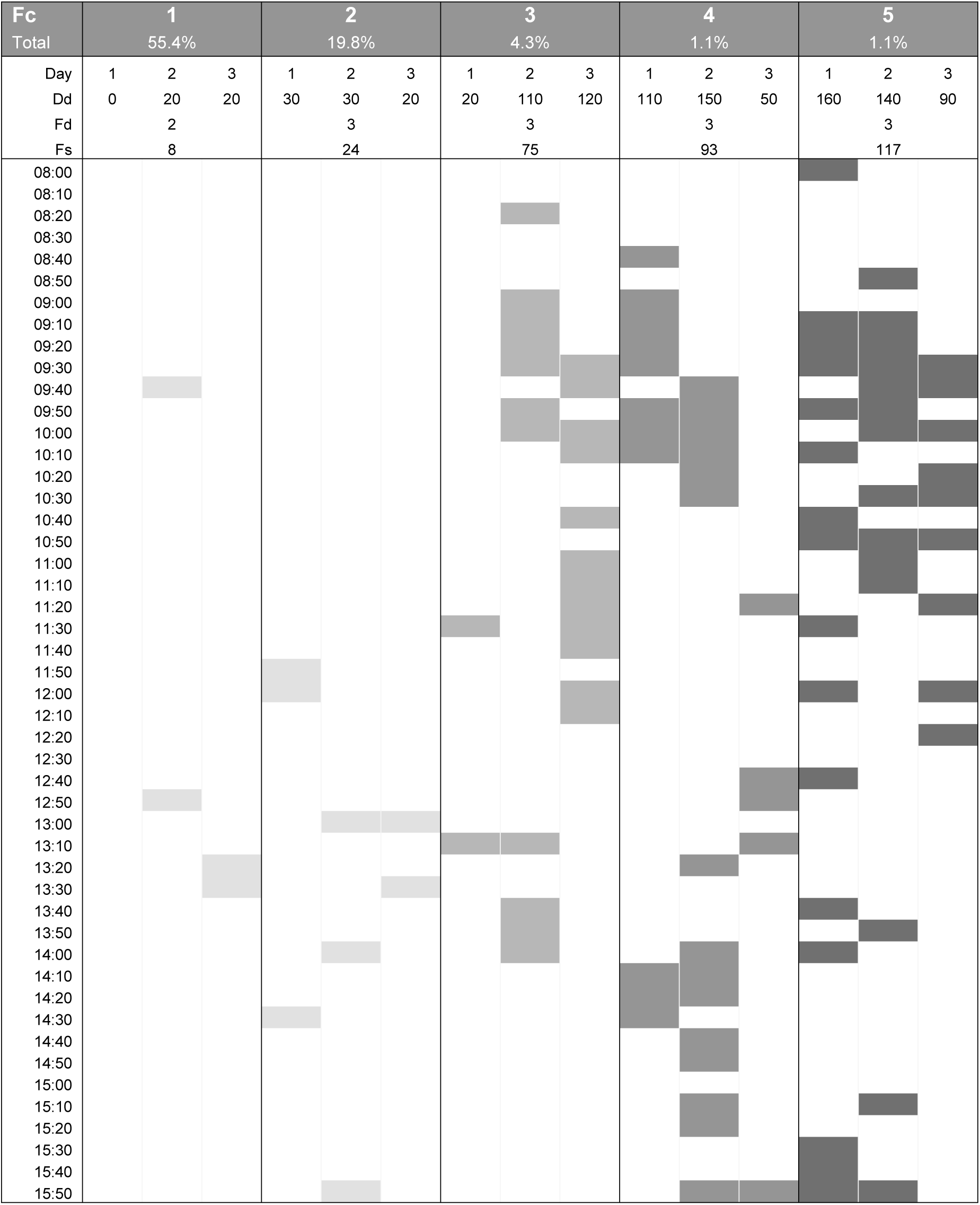
A graphic representation of selected birds’ recorded foraging activity, demonstrating the relationship between Foraging score (Fs), Daily duration (Dd), and Frequency-days (Fd). The relative frequency (over all birds in the study) of each Foraging class (Fc) 1 to 5 is shown. The relative frequency of Class 0, which is not shown, was 18.3%.

**Supplementary Figure 3:**
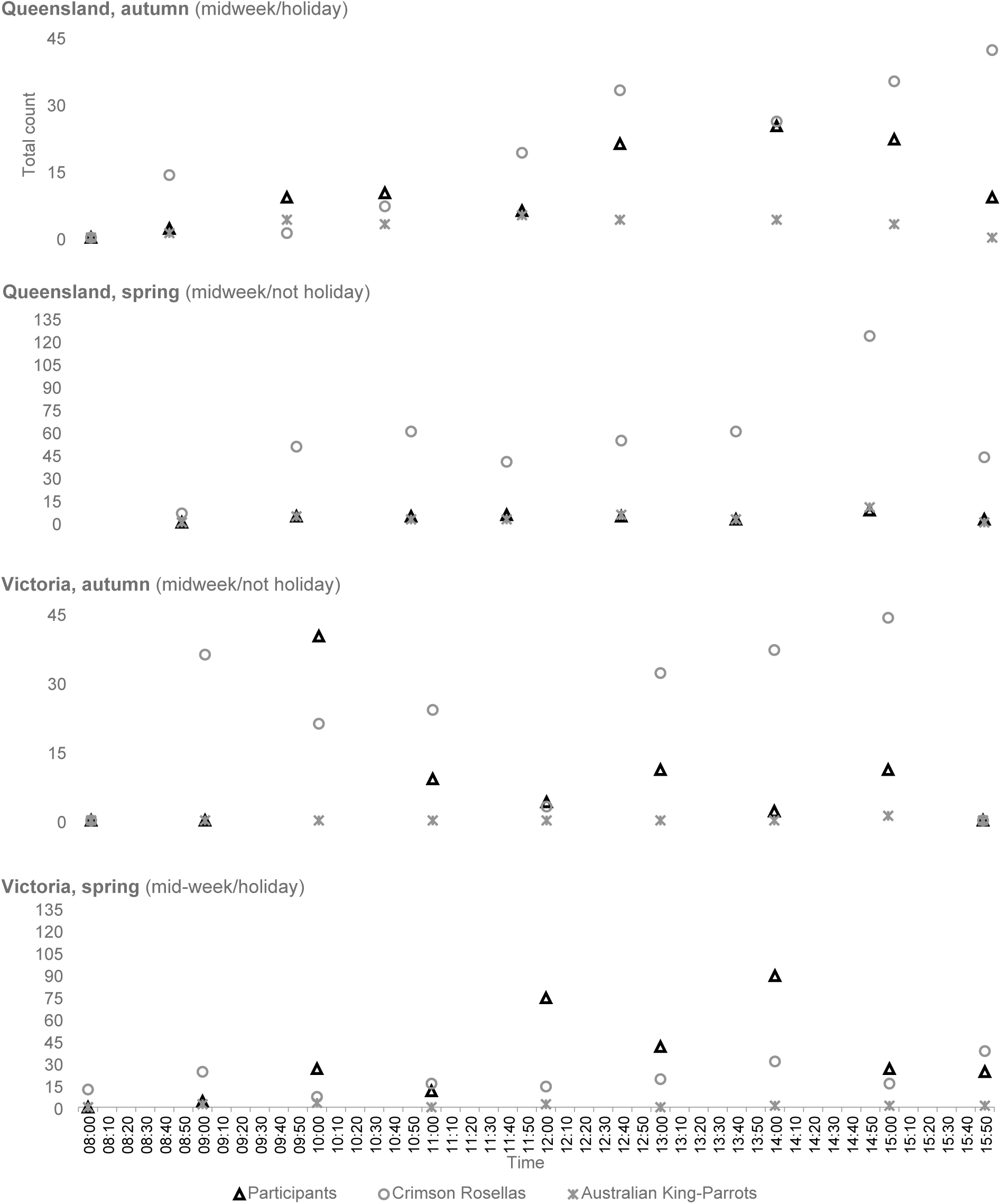
A graphic representation of total counts of Crimson Rosellas, Australian King-Parrots and participants at a Queensland and Victorian wild bird feeding site during one 10-minute interval per hour, on the first day of the spring and autumn observation periods.

